# High-temporal resolution functional PET/MRI reveals coupling between human metabolic and hemodynamic brain response

**DOI:** 10.1101/2023.08.02.551631

**Authors:** Andreas Hahn, Murray B. Reed, Chrysoula Vraka, Godber M. Godbersen, Sebastian Klug, Arkadiusz Komorowski, Pia Falb, Lukas Nics, Tatjana Traub-Weidinger, Marcus Hacker, Rupert Lanzenberger

**Author notes:** **Correspondence to:** Andreas Hahn, Assoc.Prof. PD PhD MSc, Rupert Lanzenberger, Prof. PD MD, NeuroImaging Labs (www.meduniwien.ac.at/neuroimaging) Department of Psychiatry and Psychotherapy, Medical University of Vienna, Austria Waehringer Guertel 18-20, 1090 Vienna, Austria.

## Abstract

Positron emission tomography (PET) provides precise molecular information on physiological processes, but its low temporal resolution is a major obstacle. Consequently, we characterized the metabolic response of the human brain to working memory performance using an optimized functional PET framework at a temporal resolution of 3 seconds. Consistent with simulated kinetic modeling, we observed a constant increase in the [^18^F]FDG signal during task execution, followed by a rapid return to baseline after stimulation ceased. The simultaneous acquisition of BOLD fMRI revealed that the temporal coupling between hemodynamic and metabolic signals in the primary motor cortex was related to individual behavioral performance during working memory. Furthermore, task-induced BOLD deactivations in the posteromedial default mode network were accompanied by distinct temporal patterns in glucose metabolism, which depended on the task-positive network metabolic demands. In sum, the proposed approach enables the advancement from parallel to truly synchronized investigation of metabolic and hemodynamic responses during cognitive processing.

## INTRODUCTION

The human brain has the ability to process a vast amount of information at incredibly high temporal scales, largely defined by the underlying neuronal architecture (1–3). Therefore, characterizing these biological processes at a high temporal resolution is key to strengthen our understanding of brain function. However, current non-invasive approaches are often limited in either their temporal or spatial domain (4, 5). In the worst case scenario, measurement parameters can only be obtained after the actual stimulation and the corresponding neuronal activity have ceased (6).

One of the most commonly used techniques is electroencephalography (EEG), which provides the highest temporal resolution in the millisecond range. However, EEG has limited spatial resolution, which is why blood-oxygen level dependent (BOLD) fMRI often represents the method of choice to assess human brain function *in vivo* with a good compromise between temporal (seconds) and spatial resolution (mm). As a drawback, the BOLD signal can only be indirectly related to neuronal activation (7) since it reflects a composite of blood flow, volume and oxygenation (6). In contrast, positron emission tomography (PET) has a spatial resolution comparable to that of fMRI and provides unmatched molecular sensitivity and specificity. However, standard PET imaging is often limited to snapshots of the underlying molecular processes in the range of minutes to hours.

The recent introduction of functional PET (fPET) imaging offers a solution to this problem through the constant administration of the radioligand and repeated stimulation, similar to fMRI designs (8, 9). By using the radiolabeled glucose analogue [^18^F]FDG, it enables the acquisition of baseline metabolism and task-specific changes in a single PET scan. Previous work with simultaneous assessment of fPET and fMRI has characterized the spatial agreement as well as unexpected differences between stimulation-induced changes in BOLD contrast and glucose metabolism by either acknowledging the different timescales of the signals (10) or by using a hierarchical design (11, 12). Although the temporal resolution of single fPET time frames has been improved from minutes to 16 s (13) or even 6-12 s (14), this still impedes a direct comparison between BOLD and metabolic signals.

To overcome these issues we combined multiple recent advances in PET imaging. These include i) fPET with a bolus plus constant infusion protocol to increase the signal-to-noise ratio (SNR) of the metabolic signal (14). ii) A novel dynamic filtering technique (15) with a sliding window was implemented, which enhances task-specific effects and enables reconstruction of high-temporal resolution fPET frames. iii) Finally, a truly synchronized acquisition of [^18^F]FDG fPET and BOLD fMRI data was employed during an established working memory paradigm. Notably, these developments are readily applicable to conventional PET scanner systems, thus enabling a widespread implementation without specific or costly hardware requirements. As a secondary aim, we evaluated the performance of a new-generation PET/CT scanner with time-of-flight imaging compared to an older, more widely available scanner in the context of fPET imaging.

## RESULTS

[^18^F]FDG fPET and BOLD fMRI data were acquired from 35 healthy participants during performance of the established n-back working memory task (16). Of those, 19 underwent simultaneous PET/MRI. Data of the remaining participants were acquired with a new-generation PET/CT scanner and a separate MRI system. fPET data was reconstructed at three different temporal resolutions (3, 6 and 12 s) to investigate the complementary nature between BOLD signal and glucose metabolism at different levels. First, brain regions with consistent task-specific increases between the two modalities were identified. Moving to the temporal domain, the time course of each modality was then analyzed to assess the feasibility of high-temporal resolution fPET. This was complemented by simulations of task-induced changes of the [^18^F]FDG signal across a range of variations in rate constants. Next, the individual association between hemodynamic and metabolic signals was investigated and related to working memory task performance in the simultaneous PET/MRI sample. Finally, we assessed the value of time domain analyses with respect to a recent spatial dissociation between the BOLD signal and metabolic demands in the posteromedial default mode network (12) as well as the corresponding influence of task-positive networks (17).

### Spatial domain

Statistical conjunction analysis was used to assess whether working memory performance induces similar spatial activations between metabolic and BOLD signal changes. Increases across both signals were observed in the right dorsolateral prefrontal cortex, the left primary motor cortex as well as the intraparietal sulcus and anterior insula bilaterally during the 2-back condition (p<0.05 FWE corrected). This activation pattern was robust for all temporal resolutions of 3, 6 and 12 s with Dice coefficients between 0.71 and 0.78 (Fig 1). Across these regions the average increase was K_i_ = 0.011 ± 0.005 ml/cm^3^/min and CMRGlu = 6.74 ± 2.78 µmol/100g/min, which corresponds to a change of 27.15 ± 9.91 %.

**Figure 1:**
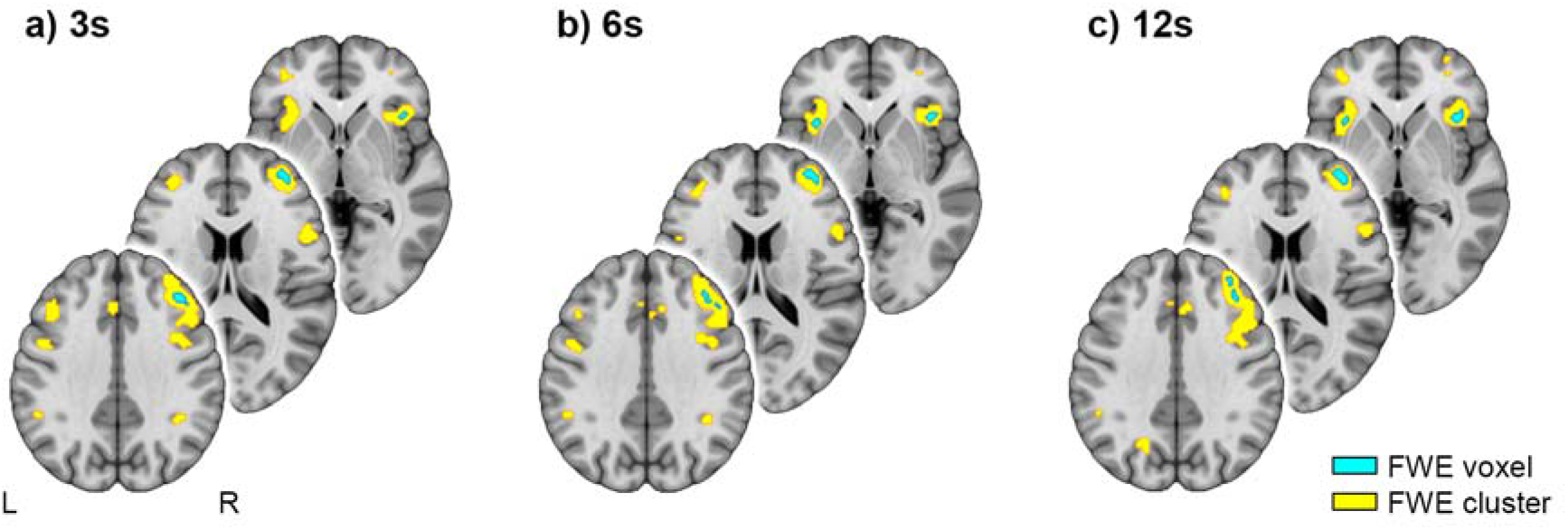
Spatial correspondence of BOLD fMRI changes and [^18^F]FDG fPET glucose metabolism induced by the 2-back task condition. Statistical conjunction analysis between the two signals yields robust task effects in several brain regions involved in working memory (16), such as the right dorsolateral prefrontal cortex, left primary motor cortex and bilateral anterior insula (blue: p<0.05 FWE-corrected voxel level, yellow: p<0.05 FWE-corrected cluster level following p<0.001 uncorrected voxel level). The effects were stable for different temporal resolutions of the fPET data, namely 3s (a), 6s (b) and 12s (c). Dice coefficients between the activation patterns obtained with the three temporal resolutions were between 0.71 and 0.78. Voxel-level corrected regions of the 3s data (i.e., blue regions in a) were used for subsequent temporal assessment. MNI coordinates for slices are z = −1, 17 and 35 mm. Left is left.

### Temporal domain

The time courses of the [^18^F]FDG metabolic and BOLD hemodynamic signals were then extracted from significant brain regions (Fig 1, p<0.05 FWE corrected voxel level) and averaged across all 2-back task blocks and participants. As expected, the BOLD signal changes exhibited a slightly delayed plateau with respect to task performance (Fig 2a). The high-temporal resolution [^18^F]FDG signal showed a continuous increase during task execution due to the irreversible binding of the radioligand. Interestingly, also the metabolic signal returned to baseline within 10 s, which was most visible for the 3 s reconstruction (Fig 2b). Therefore, the highest temporal resolution was used for all subsequent computations, further enabling direct comparison with the BOLD signal. Additional simulations were carried out to assess the temporal dynamics of brain metabolism in response to task-induced changes in the corresponding rate constants of the two-tissue compartment model. The simulated time courses showed a remarkable match with the observed task response (Fig 2c). Specifically, simulations yielded a linear increase of the signal, which returned to baseline after stimulation ceased. The simulation results were robust across a physiologically plausible range of different kinetics, which included changes in K_i_ of 12, 22 and 33%, elicited by variation of K_1_, k_2_ and k_3_ between 5 and 15% (Fig 2d). The different simulations only changed the amplitude of the time course, but not the overall shape. This was also the case when using rate constants from previous work (8).

**Figure 2:**
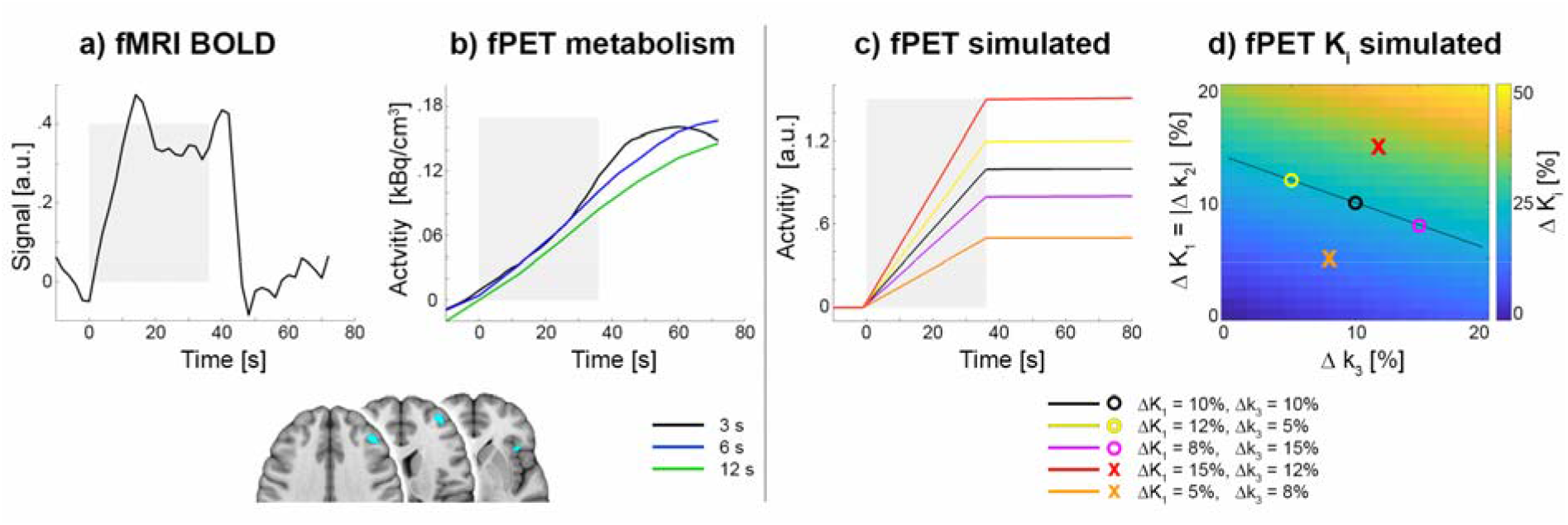
Concurrent temporal changes in BOLD fMRI and [^18^F]FDG fPET signals during working memory performance across all active regions. a) Task-induced changes in the BOLD signal show a plateau of activation as expected from the block design. b) Task-specific time course of [^18^F]FDG glucose metabolism resulted in a constant increase during task performance (due to the irreversible binding of the radiotracer) and rapid return to baseline afterwards (i.e., a steady activity level). The time course was generally similar for the different temporal resolutions of 3s (black), 6s (blue) and 12s (green), but the 3s data showed highest ability to reconstruct the return to baseline. c) Simulation of task-related changes in the [^18^F]FDG signal was in close agreement with the shape of the experimental observation in b across different variations in rate constants. Colors of lines match those of symbols in d. d) Matrix of changes in K_i_ induced by variation of rate constants K_1_, k_2_ and k_3_ from 0 to 20%. Changes in rate constants were chosen to yield physiological increases in the net influx constant K_i_ around 22% (black line as well as black, yellow and magenta circles) as observed in this and previous work (9). In addition, rate constants yielding K_i_ = 12 and 33% (i.e., orthogonal to the expected 22%-line) were investigated (orange and red x, respectively). For all simulations k_2_ = -K_1_. Gray areas in a-c indicate the time of the task performance. Data in a and b represent the average across participants, 2-back task blocks and regions obtained from the statistical conjunction analysis (blue regions in brain slices, p<0.05 FWE-corrected voxel level, identical to Fig. 1a). Data in b and c show task effects after subtraction of baseline metabolism.

Next, we investigated the individual association between metabolic and hemodynamic signals and their relationship with task performance for the sample that underwent truly simultaneous PET/MR imaging (n=17). Across all task regions, we observed a moderate agreement between both modalities (|r| = 0.31 ± 0.19). Moreover, the association of BOLD and metabolic signals in the left primary motor cortex correlated with the reaction time of correct 2-back button presses, except for one outlier whose reaction time was more than 3 standard deviations higher than the group average (rho = −0.69, p = 0.02 corrected, time window = −12 to +36 s with respect to task performance, Fig. 3d). The correlation with task performance was independent of the time windows (−12 to +24 s, −6 to +12 s and −6 to +6 s) used for the multimodal signal correlations with rho = −0.68…-0.65 (p = 0.024…0.043 corrected). The corresponding time series are exemplarily shown in figure 3a-c, indicating a similar temporal profile of the metabolic and BOLD signals with individual delays. Importantly, the association between the two signals was not driven by motion artifacts (rho = −0.15, p = 0.57).

**Figure 3:**
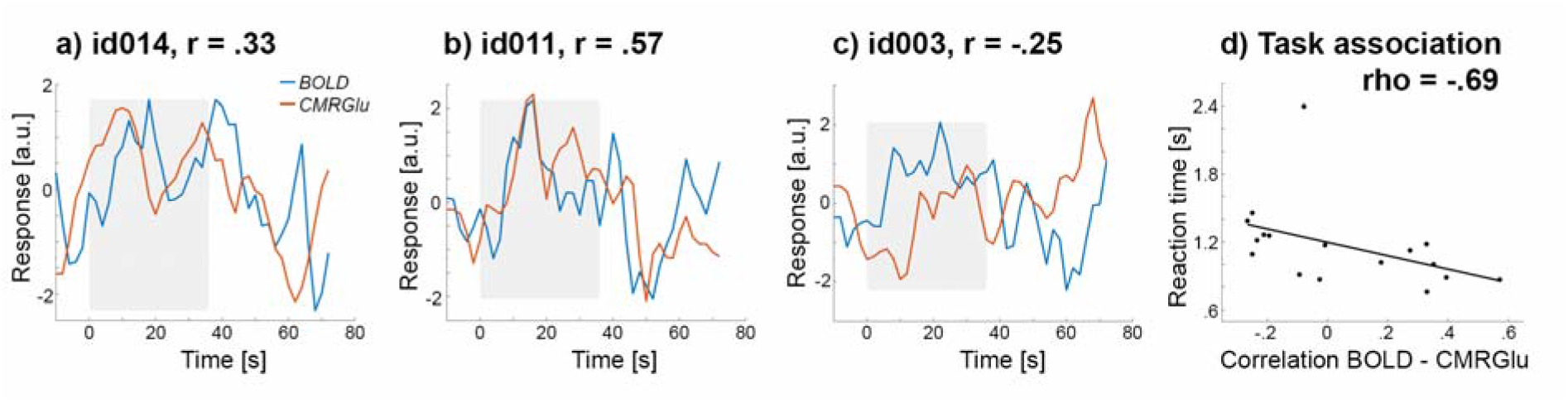
Temporal associations between signals of fMRI BOLD (blue) and fPET glucose metabolism (CMRGlu, red) in the left primary motor cortex. a-c depict three representative examples of individual time courses of the BOLD signal and [^18^F]FDG glucose metabolism, averaged across the 2-back working memory task blocks. For CMRGlu, the first derivative is shown to account for the irreversible accumulation of the radiotracer. The three time courses show that a) the metabolic signal may either precede the BOLD signal, b) both signal changes occur simultaneously or c) that the BOLD signal precedes that of CMRGlu. Data in a-c were z-transformed to account for different amplitudes of the signal. Gray areas indicate the time of the task performance. d) The individual temporal correlation between the CMRGlu and BOLD signals was associated with the corresponding response time of correct button presses during the 2-back working memory task (rho=-0.69, p=0.02 corrected).

Finally, we investigated the temporal domain in relation to previous findings of a spatial dissociation between BOLD changes and glucose metabolism in the posteromedial default mode network (DMN) during working memory (12). Consistent with those findings we observed a negative BOLD response in the posterior cingulate cortex (PCC) during the 2-back working memory execution. However, no change in metabolic demands was found when averaged across the entire sample (Fig. 4b, p<0.05 FWE-corrected cluster level following p<0.001 uncorrected voxel level). This dissociation was also evident when using the PCC as region of interest (BOLD beta = −0.93 ± 0.29, p < 10^−17^; K_i_ = 0.0006 ± 0.0072 ml/cm^3^/min, p=0.6). Given the corresponding metabolic influence of frontoparietal (FPN) and dorsal attention networks (DAN) on the PCC (17), participants were split into quartiles according to the difference in task-specific K_i_ between the FPN and DAN (Fig. 4a). This resulted in three distinct groups with the metabolism of FPN < DAN (low, 25% of participants), FPN = DAN (balanced, 50%) and FPN > DAN (high, 25%). Evaluating the temporal profile of the PCC between these groups revealed an interaction effect (p = 0.0016, Fig. 4c), with significant post-hoc differences between high vs. low participants (p < 10^−5^) and low vs. balanced (p = 0.025). In detail, the low group (i.e., with lower metabolism in FPN than DAN) was characterized by a constant decrease in PCC metabolism during task execution, followed by an increase afterwards. In contrast, the high group only exhibited a delayed increase in the metabolic signal, whereas the balanced group showed almost no variation in their metabolic response. Again, the temporal difference in the metabolic profile between the three groups was consistent across all time windows (p = 0.0005…0.040). However, the BOLD time series of the PCC showed robust decreases for all three groups, without any significant difference (Fig. 4d, group-by-time interaction p = 0.9).

**Figure 4:**
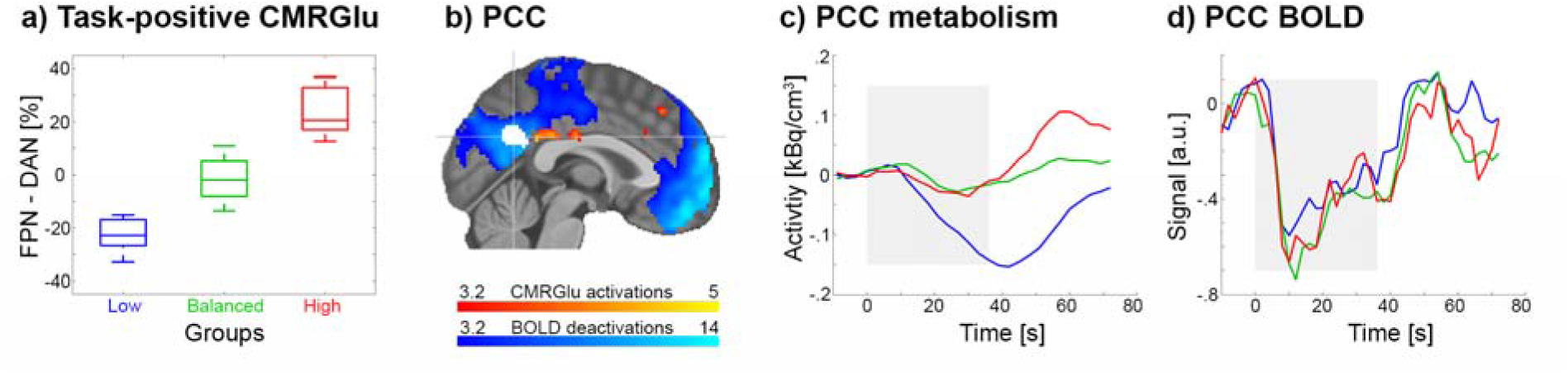
fPET metabolic and fMRI BOLD signals in the posterior cingulate cortex (PCC) during the 2-back working memory task. a) Participants were split into three groups according to significant changes in CMRGlu in the frontoparietal (FPN) and dorsal attention network (DAN, p<0.05 FWE-corrected voxel level, Fig. 1a) as these networks drive CMRGlu changes in the PCC (17). By definition, the quartile split separates 50% of participants with similar changes in these networks (green, balanced group), while 25% had lower task-specific CMRGlu in FPN than DAN (blue, low) and 25% had higher CMRGlu in FPN than DAN (red, high). b) Task-specific decreases in the BOLD signal (blue) and increases in CMRGlu (red) during working memory performance (p<0.05 FWE-corrected cluster level following p<0.001 uncorrected voxel level). Replicating previous work (12), BOLD decreases where pronounced in the PCC, while CMRGlu did not change significantly. The white area indicated by the cross-hair represents the PCC cluster of our recent study (17), which was used for c and d. c) The time course of [^18^F]FDG glucose metabolism in the PCC was significantly different between the three groups (group*time interaction p=0.0016). Participants with lower CMRGlu in FPN than DAN exhibited a decrease in PCC CMRGlu (blue) during working memory, while participants with higher CMRGlu in FPN than DAN showed a delayed positive deflection in the metabolic signal. d) BOLD signals in the PCC showed similar decreases during the task for all three groups (interaction p=0.9). Colors in a, c and d represent the same groups of participants. Boxplots in a indicate median values (center line), upper and lower quartiles (box limits) and 1.5*interquartile range (whiskers). Data in c show task effects after subtraction of baseline metabolism. Gray areas indicate the time of the task performance.

### Comparison of PET systems

In the temporal domain, there was no significant difference in the ability to identify task-specific changes in glucose metabolism between the scanners (PET/MR t = 3.13±0.98, PET/CT t = 3.71±1.49, p=0.17). Similarly, K_i_ percent signal change of regions involved in the task (Fig. 1a, p<0.05 FWE-corrected voxel level) was not significantly different between the scanners (PET/MR = 29.79±10.22 %, PET/CT = 25.54 ± 10.87%, p=0.24).

## DISCUSSION

In this work we employed high temporal resolution fPET with 3 s to investigate the dynamics of glucose metabolism during working memory performance. The metabolic response of the [^18^F]FDG signal was characterized by a constant increase during task execution and followed by a rapid return to baseline, which was consistent with simulations. Using fully synchronized fPET/fMRI we further demonstrate the feasibility to assess the individual coupling of the hemodynamic and metabolic response, whose effect in the primary motor cortex was related to working memory reaction time. Finally, analysis of PCC signals revealed that only specific subgroups exhibited decreased metabolism during the task while others showed an increased response, which was dependent on the corresponding task-positive metabolic demands. Together, these findings highlight the value of assessing brain energy metabolism in the temporal domain, which provides unique information not accessible to conventional PET imaging.

Regardless of the reconstructed frame length (3, 6 and 12 s), we observed spatially robust activations induced by working memory performance which were in line with a meta-analysis of BOLD signal changes (16). Moreover, the average increase in glucose metabolism was consistent with previous work using lower, more typical, temporal resolutions of 30 to 60 s, both in terms of absolute units of K_i_ and CMRGlu as well as percent signal changes (9, 10). This indicates that the lower SNR inherent to shorter time frames is compensated by the increased number of data points for the general linear model estimation, with the additional benefit to investigate fast metabolic changes. Altogether, these observations provide confidence for the validity to quantify task-induced changes in glucose metabolism at a high temporal resolution of 3 s. Compared to our previous work (14), this is now possible through further improvement in the protocol, specifically the combination of tracer administration as bolus + constant infusion and the novel implementation of the dynamic filter. In contrast, the use of constant infusion only (i.e., without an initial bolus) (9, 12, 13) does not seem to provide sufficient SNR to reconstruct data with such a high temporal resolution (see e.g., figure 2B in (18)). Regarding the different PET systems, the new-generation PET/CT scanner with time-of-flight imaging showed similar performance compared to the PET/MR. This may be related to the similar sensitivity of the two systems (16.4 kcps/MBq (19) and 15.0 kcps/MBq (20), respectively) and the fact that the benefit of time-of-flight is less pronounced for smaller volumes such as the head (21), which improved the spatial SNR but had little to no effect on the temporal resolution.

Our improved protocol allowed us to depict the corresponding time course of the activated brain regions in unprecedented detail. This was characterized by fast changes during cognitive performance, namely an expected constant increase in the [^18^F]FDG signal during the task but also a rapid return to baseline within 10 s once stimulation ceased. These fast metabolic changes are supported by animal studies. Specifically, almost immediate differences in extracellular availability of ATP (22) and glucose (23, 24) upon neuronal activation have been reported across different animals and approaches. Although these methods differ with respect to the return to baseline, two of these studies also matched our work in this respect with 12 ± 3 s (24) and a range between 10-100 s (22). As these differences are likely related to the specific approach and species, the exact metabolic response obtained with [^18^F]FDG as well as its variation across participants and paradigms need to be determined in future studies. Nevertheless, the time course of the task-induced metabolic signal was supported by previous (25) and our own simulations. In contrast to other work simulating slow metabolic changes due to variation of k_3_ only (8, 12), we employed a complete simulation of the irreversible two-tissue compartment model with changes in all three relevant rate constants. In particular, the variation of K_1_ is feasible as this parameter is closely related to cerebral blood flow (CBF). CBF in turn is well known to change with neuronal activation and also represents a major driver of the BOLD signal (26, 27). Furthermore, glucose as well as [^18^F]FDG are subject to transport across the blood brain barrier (BBB) that is facilitated by the glucose transporter 1 (GLUT1) carrier protein. In this transport system, however, influx from blood plasma to the brain (K_1_) will occupy the carrier protein and thus block its availability for the reverse transport (k_2_), i.e., influx and efflux rate constants are inversely related and affect each other (28–31). Although k_3_ most closely resembles hexokinase activity and the first step in glycolysis, glucose transport across the BBB reflected by K_1_ and k_2_ represents an essential (but often neglected) aspect to meet increased metabolic demands of neuronal activation (32–34). Furthermore, all three rate constants contribute to the [^18^F]FDG signal and the final quantified CMRGlu by definition, thus, supporting their combined variation in the model (25) and the resulting temporal profile of glucose metabolism.

The coupling between the BOLD signal and CMRGlu has previously been shown to increase during task execution at the spatial level (10). The current work extends this finding to the time domain, where stronger temporal coupling between metabolic and BOLD time series was associated with faster working memory performance. Both of these signals are related to neuronal activity via different yet intertwined mechanisms (33). The BOLD contrast results from disproportional increases in oxygen consumption (CMRO_2_) and CBF (26, 27). The rise in CBF is related to glutamate release during neuronal activation (35, 36) to maintain the supply of nutrients, termed neurovascular coupling. On the other hand, neurometabolic coupling links glutamate release to an increase in glucose consumption to meet the energy demands for the reversal of ion gradients (37–39). These changes in CBF, CMRO_2_, CMRGlu and thus also the neurovascular and the neurometabolic responses have previously been considered as parallel processes driven by neuronal activation, instead of being a serial connection of events (24, 40). This may explain the observed delay between metabolic and BOLD signals in the primary motor cortex, which varied across participants. We therefore put forward the hypothesis that the individual differences in the coupling between BOLD and [^18^F]FDG signals potentially informs us about the integration of the metabolic and hemodynamic interplay in neuronal processing. Possible underlying reasons for these differences could be inter-individual differences in the hemodynamic response (41) as well as regional variability in the neurovascular unit (42, 43) and BBB glucose transport (44), which probably further translates to individual differences across participants and may also be present in a similar fashion for neurometabolic coupling. On the other hand, dissociation between the signals may provide promising information about alterations in different brain disorders. For instance, in healthy aging increased BOLD-derived activations were not matched by glucose metabolism (45). Moreover, patients with Alzheimer’s disease suffer from both impaired neurovascular coupling (46) and widespread decreased metabolism (47).

Our analysis in the temporal domain also provides further insight into the divergence between glucose metabolism and BOLD deactivations in the PCC (12). In line with our previous work in the spatial domain (17), we observed that the metabolic time course is dependent on energy demands of the corresponding task-positive networks involved in the cognitive process. More specifically, individuals with lower CMRGlu in FPN than DAN exhibited consistent negative deflections in both metabolic and BOLD time series. In contrast, higher CMRGlu in FPN than DAN did not lead to relevant changes in the metabolic signal during working memory. Since BOLD signal decreases without affecting metabolism seem unlikely (39), we speculate that metabolically expensive suppression (12, 17) and downregulation of neuronal signaling (presumably mediated by GABA and glutamate, respectively) (17) occur simultaneously, thus resulting in BOLD deactivations but a net metabolism change around zero. Interestingly, right after task execution both groups showed an increase in glucose metabolism. This may indicate that the actual return of the BOLD signal to baseline also requires energy. The effect may potentially be related to increased glutamate signaling, which leads to increases in both glucose metabolism and the BOLD signal (see above).

Since the BOLD signal in the PCC showed a negative response throughout, we assume that the ratio between CBF and CMRO_2_ is also similar for all participants. As a consequence, the variation of glucose metabolism across individuals and time further implies that also the oxygen-to-glucose index (OGI) differs in the same manner. In particular, those individuals without a decreased metabolic response (and thus higher CMRGlu in FPN than DAN) would exhibit a decreased OGI in the PCC during task performance, indicating that energy is supplied by aerobic glycolysis (48). This is supported by recent work suggesting that immediate energy demands are preferably met by aerobic glycolysis (49). However, to verify our hypothesis acquisition of the actual CMRO_2_ time series is required to fully understand the mechanisms of neurometabolic coupling (40).

In conclusion, being the first study that employs such a high temporal resolution of fPET data, our work does not claim to provide definitive answers to the observed phenomena. Rather, the approach aims to spark future discussions on the relationship between hemodynamic and metabolic signals of the human brain. Further effort is required to identify the underlying mechanisms of the coupling, particularly when the signals diverge, the delay between BOLD and metabolic time series as well as potential dissociations in brain disorders. Nevertheless, the direct comparison of these signals in the temporal domain offers numerous possibilities to investigate these aspects in health and disease, thereby introducing a previously unseen perspective to our understanding of brain function.

## METHODS

Data acquisition, blood sampling, quantification of CMRGlu (10, 14, 50) and estimation of BOLD signal changes (51) were conducted analogous to our previous work, unless otherwise specified.

### Experimental design

In this cross-sectional study participants underwent either a simultaneous fPET/fMRI (n=19) or separate fPET and fMRI scans (n=16, 5.2 ± 5.8 days between scans) with the radiotracer [^18^F]FDG. During each of the scans an established working memory task was completed in a conventional block design. fPET started with an initial resting period of 8 min, which was followed by 12 min of task performance. The initial baseline was omitted for the MRI-only scan, but the task was otherwise identical. MRI acquisitions included a T1-weighted structural image and a BOLD sequence.

### Cognitive task

The established n-back working memory task (16) was implemented in Psychtoolbox for Matlab. In this task letters were presented on a screen for 0.5 s, followed by an interstimulus interval of 1.9 s. In the 0-back control condition, a button press was required when the letter “X” appeared on the screen. For the 2-back condition, a button press was required when the current letter was the same as the one shown two letters before. Before each task block the corresponding instructions were shown for 2 s. The two conditions were shown in blocks of 36 s, each comprising 15 stimuli and 5 button presses (pseudo-randomized order). Resting periods were included at the beginning of the task (14 s), after each instruction (2 s) and after each task block (32 s). For each condition 5 blocks were presented in pseudo-randomized order, resulting in a total task duration of 12 min. The letters used in the task were phonologically similar in the German alphabet to avoid learning strategies (52, 53). Small as well as capital letters were used equally (b, c, d, g, p, t, w). During all periods of rest, participants were instructed to look at a crosshair, relax and not to focus on anything in particular. Before data acquisition, participants completed a training session of each task condition inside the scanner to familiarize themselves with the task and the controls.

### Participants

For this study 42 participants were initially recruited, and data from 35 were used in the final analysis (mean age ± sd = 24.5 ± 4.4 years, 19 women). Dropout reasons included: blood sampling failures (n=3), problems with the tracer administration (n=3) and gross anatomical abnormalities (n=1). Additionally, BOLD data could not be used for two participants due to technical reasons, blood glucose levels were not available for one subject and working memory performance data for another one due to failure of the response box. As no studies with such a high temporal resolution of fPET data are available (previously 6-12 s (14) and 16 s (13)), we aimed for a sample size that exceeds previous work with robust task activation by at least 50%. Each participant underwent an initial screening visit, where general health was ensured. Routine medical examinations included blood tests, electrocardiography, neurological testing and the structural clinical interview for DSM-IV. Female participants underwent pregnancy tests at the screening visit and before the PET and/or MRI scans. Further exclusion criteria were current and previous severe somatic, neurological and psychiatric conditions, substance abuse or medication, pregnancy or breastfeeding, MRI contraindications and previous study-related radiation exposure. At the screening visit, all participants provided written informed consent after detailed explanation of the study protocol. Participants were reimbursed and insured during the study. The study was approved by the ethics committee of the Medical University of Vienna (ethics numbers 1479/2015 and 2054/2020) and all procedures were carried out according to the Declaration of Helsinki.

### Data acquisition

Simultaneous acquisition of fPET and fMRI data was done with a mMR scanner system. Separate acquisition was carried out with a Biograph Vision PET/CT and a Prisma MRI scanner (all Siemens Healthineers, Erlangen, Germany).

Participants had to fast at least 5.5 hours before the start of the fPET scan (except for unsweetened water) (54). The radiotracer [^18^F]FDG was administered in a bolus (816 ml/h for 1 min) plus constant infusion protocol (114.9 ml/h for 19 min, total of 50 ml) for the entire fPET scan using a perfusion pump (Syramed µSP6000, Arcomed, Regensdorf, Switzerland), which was kept in an MR-shield (UniQUE, Arcomed).

With the PET/MR scanner, BOLD fMRI was acquired simultaneously with fPET (EPI sequence, TE/TR = 30/2000 ms, flip angle = 90°, matrix size = 80×80, 34 slices, voxel size = 2.5×2.5×3.325 mm, 12 min). Furthermore, a structural MRI was recorded before fPET acquisition, to rule out gross anatomical abnormalities and for spatial normalization (T1-weighted MPRAGE sequence, TE/TR = 4.21/2200 ms, TI = 900 ms, flip angle = 9°, matrix size = 240×256, 160 slices, voxel size 1×1×1.1 mm, 7.72 min).

For the separate MRI acquisition, corresponding imaging data were acquired (EPI sequence: TE/TR = 30/2050 ms, flip angle = 78°, matrix size = 100×100, 35 slices, voxel size = 2.1×2.1×3.5 mm, 12 min; T1-weighted MPRAGE sequence: TE/TR = 2.91/2000 ms, TI = 900 ms, flip angle = 9°, matrix size = 240×256, 192 slices, voxel size 1×1×1 mm, 8 min).

### Blood sampling

Before the fPET scans, blood glucose levels were determined (Glu_plasma_). During the fPET scans, arterial blood samples were taken manually from the left radial artery. This included sampling every 20 s for the first three minutes, and thereafter at 4, 5, 8, 12, 16 and 20 min. To avoid interference with task performance, samples after 5 min were taken remotely (outside 20 mT line) with a VAMP system. Whole-blood activity was measured in a gamma counter (Wizard^2^, Perkin Elmer). Samples obtained after 3 min were centrifuged and the plasma activity was measured. Whole-blood data were linearly interpolated to match fPET frames and multiplied with the average plasma-to-whole-blood ratio to obtain the arterial input function.

### fPET preprocessing

fPET data were corrected for attenuation using a database approach (55) and reconstructed to frames of 3, 6 and 12 s (matrix size = 344×344, 127 slices). Preprocessing was carried out in SPM12 (https://www.fil.ion.ucl.ac.uk/spm/), which included motion correction (quality=1, register to mean) and spatial normalization to MNI space with the T1-weighted structural image.

We implemented a novel filtering technique in Matlab to increase the low SNR inherent to high-temporal resolution PET data since short frames contain less radioactive counts. The filter represents a dynamic non-local means (NLM) filter (15) with a modified local patch selection. Generally, NLM filters create a weighted average of neighboring voxels within a given search window in the spatial and temporal domain. These types of filters have been shown to provide an improved contrast-to-noise ratio in dynamic PET images compared with other more conventional denoising approaches (56). The novel aspect implemented here is a sliding-window approach, instead of using the entire PET time course (15). This confines the denoising of each voxel both temporally and spatially, preventing the baseline [^18^F]FDG uptake to drive the signal and thereby specifically strengthening task-specific effects. The local neighborhood was selected with a sliding-window approach for each voxel using a spatiotemporal patch of 3×3×3 voxels and 18 s in a search window of 11×11×11 voxels. The weight is given by the similarity between voxels (i.e., Gaussian distance from center voxel) as in the original description (15). Data were subsequently smoothed with a 5 mm Gaussian kernel in SPM12. Together with the NLM filter, this corresponds approximately to a total kernel of 8 mm, which facilitates comparison with BOLD data and previous work.

### Cerebral metabolic rate of glucose (CMRGlu)

Quantification of CMRGlu was carried out voxel-wise. Data were masked to include only gray matter. To further improve the SNR in the temporal domain, a low pass filter was used with a cutoff frequency of 18 s, which represented half the task block duration. To separate task-specific effects from baseline metabolism, the general linear model (GLM) was used, which included one regressor for the baseline, one for each task condition and motion regressors. The baseline regressor was defined as the average fPET signal across all gray matter voxels, excluding voxels activated during the individual BOLD fMRI (2-back vs. rest, p<0.001 uncorrected) and those identified in a meta-analysis (16). This combination of activation maps ensures that task effects do not contaminate the baseline and yields an optimal model fit (14). The task regressors were defined as a ramp function with slope = 1 kBq/frame. For the motion regressors, the realignment parameters obtained from SPM12 were subject to a principal component analysis and a variable amount of regressors were included using the elbow method, to ensure that the majority of explained variance is covered. After the GLM, the Patlak plot was used for absolute quantification of glucose metabolism. This yields the influx constant K_i_, which reflects the combination of the individual rate constants K_1_-k_3_

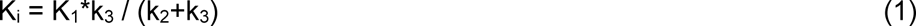

K_i_ was then converted to the cerebral metabolic rate of glucose (CMRGlu)

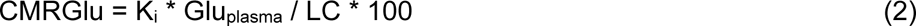

with LC being the lumped constant, which was set to 0.89 (57). This results in task-specific maps of K_i_ and CMRGlu, which were used for subsequent analysis.

### Blood-oxygen level dependent (BOLD) signal changes

BOLD data were processed in SPM12 (51). Images were corrected for slice timing effects (reference = middle slice) and head motion (quality = 1, register to mean), followed by spatial normalization to MNI space and spatial smoothing with an 8 mm Gaussian kernel. Task-related changes in the BOLD signal were computed with the GLM, including one regressor for each condition (0-back, 2-back) as well as potentially confounding signals (instructions, 24 motion parameters (58), white matter and cerebrospinal fluid). The autocorrelation option was set to FAST (59). The contrast of interest was 2-back vs. rest to facilitate comparison with fPET data.

### Statistical analyses

For statistical analyses data from n=35 participants were used for [^18^F]FDG glucose metabolism and n=33 for BOLD signal analyses (n=2 missing due to technical reasons). For correlation between the signals in the temporal domain and association with task performance, n=17 with simultaneous PET/MRI acquisition were used.

#### Spatial conjunction of task-specific neuronal activation

To assess the spatial overlap between changes in the BOLD signal and metabolic demands as induced by the 2-back working memory task, a statistical conjunction analysis was carried out in SPM12 (17). BOLD and K_i_ maps were individually z-scored and included in a one-way ANOVA with each modality representing a “group” and an additional factor accounting for the different scanners (i.e., PET/MR vs. separate PET/CT and MRI). K_i_ was used instead of CMRGlu as blood glucose data was missing for one subject. The conjunction was corrected for multiple comparisons using family-wise error rate (FWE) at the voxel level (p<0.05) and clusters with less than 5 voxels were removed. This was carried out for the three different fPET reconstructions of 3, 6 and 12 s. The Dice coefficient was used to assess the spatial agreement between the three resulting conjunction maps. For completeness and comparison with previous work (12, 17), inference was also computed at p<0.05 FWE-corrected cluster level after an initial voxel threshold of p<0.001 uncorrected.

#### Time-domain analyses

Task-specific signals were obtained as GLM residuals after removing all effects except those of the task. The resulting BOLD signal and [^18^F]FDG time series were extracted from significant voxels of the above spatial conjunction (p<0.05 FWE-corrected voxel-level). For visualization, these were averaged across 2-back blocks, participants and voxels for a time window starting 12 s before the task began until 36 s after the task (total of 84 s). Next, the BOLD and metabolic signals were correlated within each subject to evaluate the individual coupling between the two imaging modalities. These correlation values were further associated with the working memory performance, specifically the average reaction time of correct responses during the 2-back condition. This was done separately for each brain region (three clusters of the right DLPFC, left primary motor cortex, right insula, left and right intraparietal sulcus). Spearman’s rho was used to account for an outlier with substantially slower reaction time (> 3*sd of the group) and corrected for the seven brain regions using the Bonferroni method. To assess the robustness of the brain-behavior association, the time window was also constrained to −12…+24 s, - 6…+12 s and −6…+6 s, followed by re-calculation of the correlation. To rule out that individual associations between metabolic and BOLD signals are driven by motion, these values were correlated with framewise displacement as obtained from the BOLD realignment parameters (60).

#### Analyses of the posterior cingulate cortex (PCC)

We further investigated a previously reported spatial dissociation of glucose metabolism and BOLD responses in the PCC (12). First, separate maps with a negative BOLD response and a positive metabolic response during the 2-back working memory task were computed (p<0.05 FWE-corrected cluster level following p<0.001 uncorrected voxel level). Specific attention was given to the PCC, as we recently showed an opposite influence of the FPN and DAN metabolism onto the PCC (17). Thus, the difference in task-specific K_i_ between FPN and DAN during the 2-back working memory condition was computed. Percent signal change was used to enable better comparison between the different networks. Based on this difference, participants were split into three groups comprising those where metabolism of FPN < DAN (lower quartile, low group), FPN = DAN (interquartile range, balanced group) and FPN > DAN (upper quartile, high group). [^18^F]FDG metabolic time series were then extracted for the PCC region (17) and the three groups, which were compared using a repeated measures ANOVA, where the interaction between groups by time served the relevant outcome parameter. Similarly, BOLD signal time courses were extracted from the PCC and compared in a repeated measures ANOVA.

#### Scanner comparison

The secondary aim was to investigate differences between PET scanners, here we compared a widely used mMR PET/MR to a new generation Vision PET/CT with time-of-flight (both Siemens) in the feasibility to compute high-temporal resolution fPET. For the subsequent analyses, data were extracted from regions with significant activation during the 2-back condition (p<0.05 FWE corrected voxel-level, Fig 1a). In the temporal domain the identifiability of task-specific changes in glucose metabolism is given by t-values of the GLM analysis, i.e., the estimated effect size divided by the standard error (14). A two-sample t-test between the scanners was performed. Next, K_i_ percent signal change was compared between the scanners, again with a two-sample t-test.

### Simulations

Simulations were carried out to assess whether the observed time course of the [^18^F]FDG metabolic signal matches theoretical calculations. The two-tissue compartment model was implemented in Matlab with an average arterial input function and rate constants as observed in our previous work at resting state (K_1_ = 0.0827 ml/cm^3^/min, k_2_ = 0.0771 min^−1^, k_3_ = 0.0629 min^−1^, k_4_ = 0 min^−1^) (14, 50). Task-specific changes in rate constants were introduced to simulate physiologically plausible increases in K_i_ (see equation 1). Previous work showed task-induced changes in K_i_ around 22% (9, 10), but for completeness also increases of 12 and 33% were introduced. These values were obtained by the following changes in rate constants i) K_1_ = 10% and k_3_ = 10%, ii) K_1_ = 12% and k_3_ = 5%, iii) K_1_ = 8% and k_3_ = 15%, iv) K_1_ = 15% and k_3_ = 12%, v) K_1_ = 5% and k_3_ = 8%. In all cases k_2_ = -K_1_ (28–31). Please see discussion for a neurophysiological rationale of these parameter variations. In addition, the simulation was repeated with baseline rate constants of previous work (K_1_ = 0.1 ml/cm^3^/min, k_2_ = 0.15 min^−1^, k_3_ = 0.08 min^−1^, k_4_ = 0 min^−1^) (8).

## ACKNOWLEDGEMENTS

We thank the graduated team members and the diploma students of the Neuroimaging Lab (NIL, head: R. Lanzenberger) as well as the clinical colleagues from the Department of Psychiatry and Psychotherapy for clinical and/or administrative support. In detail, we would like to thank S. Kasper, K. Papageorgiou, L. Silberbauer, C. Schmidt, B. Eggersdorfer, J. Unterholzner and V. Popper for medical support, L. Rischka for acquisition and analysis support, V. Ritter and C. Wotawa for subject recruitment. We are further grateful to W. Wadsak, V. Pichler, G. Karanikas, W. Langsteger and the radioligand synthesis team from the Department of Biomedical Imaging and Image-guided Therapy, Division of Nuclear Medicine for acquisition support and supervision. The scientific project was performed with the support of the Medical Imaging Cluster of the Medical University of Vienna.

## AUTHOR CONTRIBUTIONS

Study design: A.H., R.L., T. T-W., M.H.

Data acquisition: C.V., L.N., G.M.G., S.K., A.K., M.B.R., A.H.

Methods: M.B.R., A.H., P.F.

Data analysis: M.B.R., A.H.

Manuscript preparation: A.H., M.B.R., G.M.G., S.K., R.L.

All authors discussed the implications of the findings and approved the final version of the manuscript.

## DATA and CODE AVAILABILITY

Raw data will not be publicly available due to reasons of data protection. Processed data and custom code can be obtained from the corresponding author with a data-sharing agreement, approved by the departments of legal affairs and data clearing of the Medical University of Vienna.

